# Gender, race and parenthood impact academic productivity during the COVID-19 pandemic: from survey to action

**DOI:** 10.1101/2020.07.04.187583

**Authors:** Fernanda Staniscuaski, Livia Kmetzsch, Eugenia Zandonà, Fernanda Reichert, Rossana C. Soletti, Zelia M.C Ludwig, Eliade F. Lima, Adriana Neumann, Ida V.D. Schwartz, Pamela B. Mello-Carpes, Alessandra S.K. Tamajusuku, Fernanda P. Werneck, Felipe K. Ricachenevsky, Camila Infanger, Adriana Seixas, Charley C. Staats, Leticia de Oliveira

**Author notes:** Corresponding author. Fernanda Staniscuaski, **Address:** Bento Gonçalves 9500, Building 43431 Room 214 - Porto Alegre -RS - Brazil, 91501-970., **Email:**.

## Abstract

While the Coronavirus disease 2019 (COVID-19) pandemic is altering academia dynamics, those juggling remote work and domestic demands – including childcare - have already felt the impacts on productivity. Female authors are facing a decrease in papers submission rates since the beginning of the pandemic period. The reasons for this decline in women productivity need to be further investigated. Here we show the influence of gender, parenthood and race in academics productivity during the pandemic period, based on a survey answered by 3,345 Brazilian academics from various knowledge areas and research institutions. Findings revealed that male academics - especially childless ones - were the least affected group, whereas female academics, especially Black women and mothers, were the most impacted group. This scenario will leave long-term effects on the career progression of the most affected groups. The results presented here are crucial for the development of actions and policies that aim to avoid further deepening the gender gap in science. This particular situation we are facing during the pandemic demands institutional flexibility and academia should foster the discussion about actions to benefit Black scientists and academics with families in the post-pandemic scenario.

## Introduction

As COVID-19 spreads around the globe, countries are facing different degrees of lockdown and social distancing^1^. In most affected countries, schools and universities have turned the usual lectures into online classes and remote activities. Notably, the pandemic is altering many scientists’ working dynamics, especially so for the parents of young children^2^, as these scientists are thus faced with the extra challenge of accommodating remote work with domestic labor, which includes full-time child-care responsibilities. The latter are tasks predominantly performed by women^3–5^. Since the pandemic outbreak, editors from a variety of respectable scientific journals have now warned the scientific community of the increasingly fewer manuscript submissions authored by women, despite the overall increase in total submissions, driven by submissions made by male authors. The effect is even more striking from the perspective of women as first authors^6^. Therefore, it is expected that the gender gap in productivity could increase after the pandemic, but it is not clear whether mothers will be more impacted. In addition, underrepresented groups in science, such as Black female academics, who represent a very thin portion of the overall faculty population^7^, are also expected to suffer a greater impact of the pandemic-related circumstances. This impact will most likely cause effects on their career progression and overall ascension in academia.

This evidence raises an urgency to bring to light the full picture of the pandemic impact on the careers of female academics, especially the mothers and those from underrepresented groups in science. Also, the identification of the impacts in scientific communities in developing countries should be a top priority behind the design of mitigation policies aimed at building more inclusive research capacities. In order to do that, we report herein the impact of Covid-19 related social isolation on the academic productivity of scientists in Brazil, focusing on the influences of the factors gender, parenthood, and race. We collected data via an online survey broadly disseminated across Brazilian regions and research institutions over a month of social isolation period. It was answered by 3,345 scientists. For the purpose of this study, academic productivity is regarded as the ability to submit papers within schedule and to meet overall deadlines in the pandemic period. The design of the survey used was aimed at providing a comprehensive assessment of the various elements of academic productivity, relevant to the full array of knowledge areas and research institutions.

## Results

Male academics productivity, both in terms of manuscript submission (Fig. 1A) and the ability to meet deadlines (Fig. 1B), has been less affected by the pandemic circumstances than that of women. There was a significant difference between men and women for the submission of manuscripts (χ2 = 88.42, P < 0.0001) (Fig. 1A). Positive associations between women and non-submission of manuscripts, as well as between men and submission of manuscripts were observed (Fig. 1A). There was a significant difference between men and women (χ2 = 21.73, P < 0.0001) in meeting deadlines (Fig. 1B). Positive associations between women and failure to meet deadlines and between men and meeting the deadlines were observed (Fig. 1B). There was a significant effect of parenting for the submission of manuscripts (χ2 = 110.79, P < 0.0001) (Fig. 1C). There was a positive association between women with children and non-submission of manuscripts. However, no association could be observed for women without children. The proportion of men without children that submitted is higher in comparison to men with children (P < 0.01, Bonferroni PostHoc test) (Fig. 1C). Also, the proportion of women without children that submitted is higher in comparison to women with children (P < 0.01, Bonferroni PostHoc test) (Fig. 1C). There was a significant effect of parenting for meeting the deadlines (χ2 = 55.33, P < 0.0001) (Fig. 1D). Positive associations between women with children and failure to meet deadlines and between men without children and meeting the deadlines were detected. There was a significant difference (P < 0.0001, Bonferroni PostHoc comparison) comparing the proportion of women and men with children that met the deadlines (Fig. 1D). Moreover, the proportion of women without children that met the deadlines is higher than women with children (P < 0.0001 Bonferroni PostHoc comparison) (Fig. 1D).

**Figure 1.**
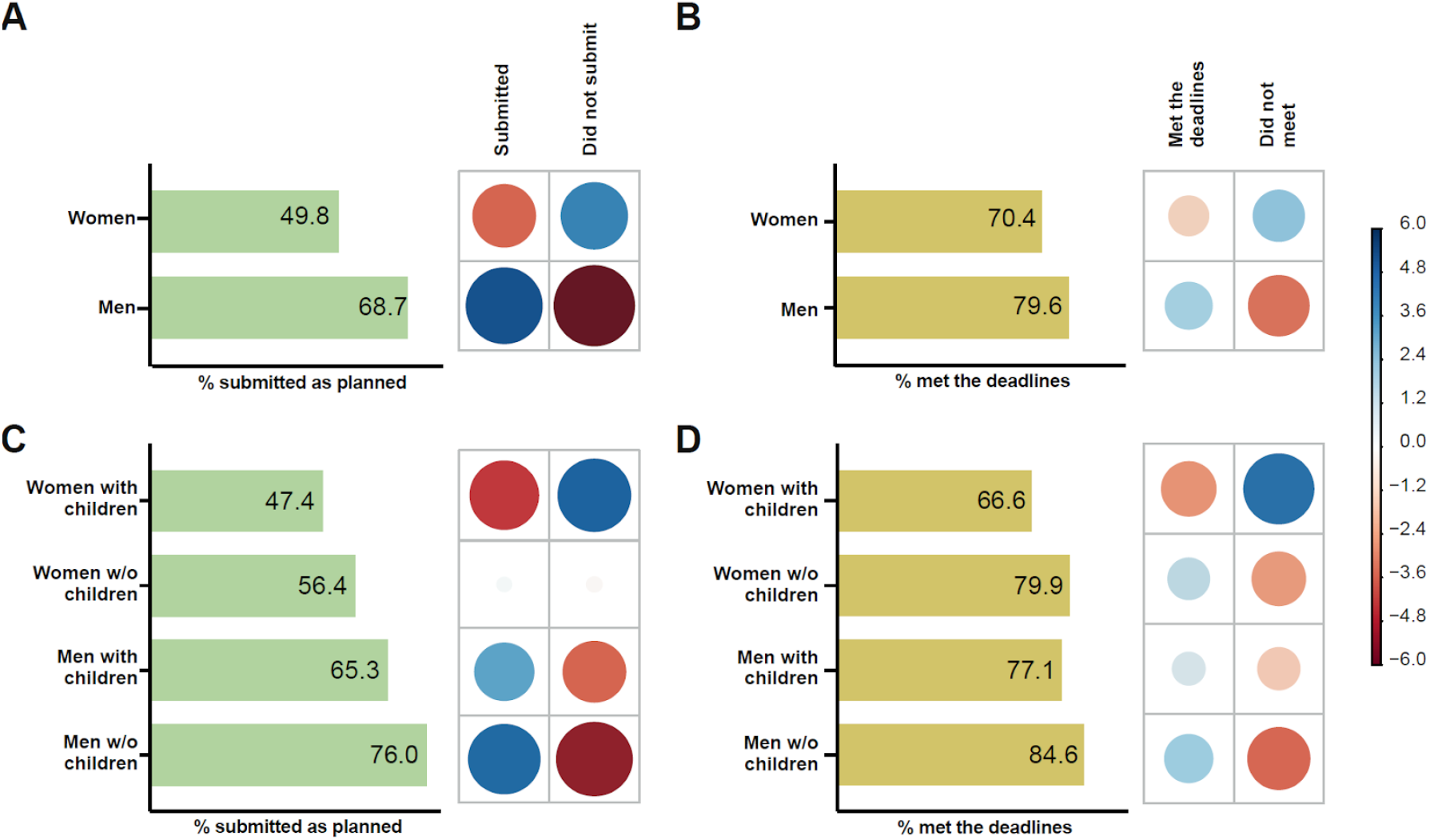
The impact of gender and parenthood on manuscript submissions (A, C) and meeting deadlines (B, D) during COVID-19 pandemic. For each figure, the graph on the left-hand side represents the percentage of respondents that submitted manuscripts as planned (A, C) or met the deadlines (B, D), while on the right-hand side the correlation plot shows Pearson’s chi-squared standardized residuals calculated for each group. Positive residuals (blue) indicate a positive correlation, whereas negative residuals (red) indicate a negative correlation. The size of the circle is proportional to the amount of the cell contribution to the χ2 score. A. Gender effect on submissions. B. Gender effect on meeting deadlines. C. Parenting effect on submissions. D. Parenting effect on meeting the deadlines.

There was no overall racial effect (Black vs White researchers) in productivity during the pandemic period for either submissions (χ2 = 2.29, p = 0.1304) nor for meeting the deadlines (χ2 = 0.06, p = 0.7956) (Fig S1). There was a significant effect of race and gender for the submission of manuscripts (χ2 = 91.01, P < 0.0001) (Fig. 2A) and for meeting the deadlines (χ2 = 21.39, P < 0.0001) (Fig. 2B). Positive associations between White men and submission of manuscripts, as well as among women, both Black and White, and non-submission of manuscripts were observed (Fig. 2A). A positive association between White women and failure to meet the deadlines was observed (Fig. 2B). A negative association between White men and failure to meet the deadlines was also detected (Fig. 2B). There was a significant difference comparing the proportion of White men and White women that met the deadlines (P < 0.0001, Bonferroni PostHoc comparison).

**Figure 2.**
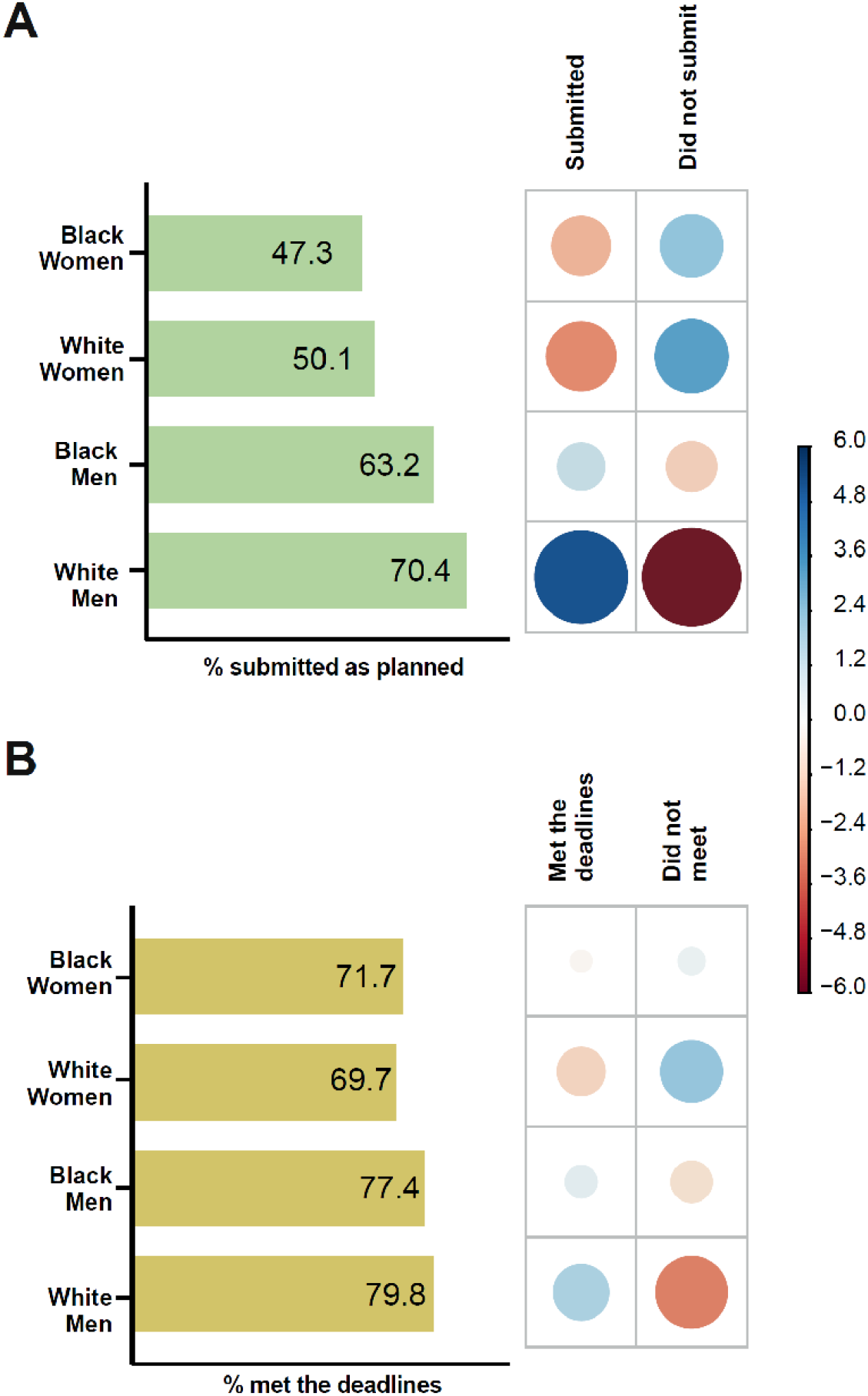
The influence of race and gender on the submission of manuscripts as planned (A) and on meeting deadlines (B) during COVID-19 pandemic. Left-hand panels show the percentage of men or women, Black or White, that submitted manuscripts as planned (A) and met the deadlines (B). The right-hand panels show the correlation plot with the Pearson’s chi-squared standardized residuals calculated for each group. The color of the circles indicates a positive correlation (blue) or negative correlation (red) and the size of the circles are proportional to the amount of the cell contribution to the χ2 score.

There was a significant difference among groups of men (Black with children, Black without children, White with children, White without children) for the submission of manuscripts (χ2 = 10.93, P < 0.05) (Fig. 3A). A negative association between White men without children and non-submission of manuscripts was detected. The proportion of White men without children that submitted manuscripts is higher than the observed for White men with children (P < 0.05, Bonferroni PostHoc test) (Fig. 3A). There was a significant difference among groups of women (Black with children, Black without children, White with children, White without children) for the submission of manuscripts (χ2 = 16.43, P < 0.001) (Fig. 3B). There was a positive association between White women without children and submission of manuscripts. The proportion of White women without children who submitted manuscripts is higher than that observed for White women with children (P < 0.01, Bonferroni PostHoc test) (Fig. 3B). There was not a significant difference between groups (Black with children, Black without children, White with children, White without children) among men (χ2 = 5.15, P = 0.1611) (Fig. 3C). There was a significant difference among groups of women (Black with children, Black without children, White with children, White without children) for meeting the deadlines (χ2 = 20.62, P < 0.01) (Fig. 3D). There was a negative association between White women without children and failure to meet the deadlines. The proportion of White women without children that met the deadlines is higher than the observed for White women with children (P < 0.001, Bonferroni PostHoc test) (Fig. 3D).

**Figure 3.**
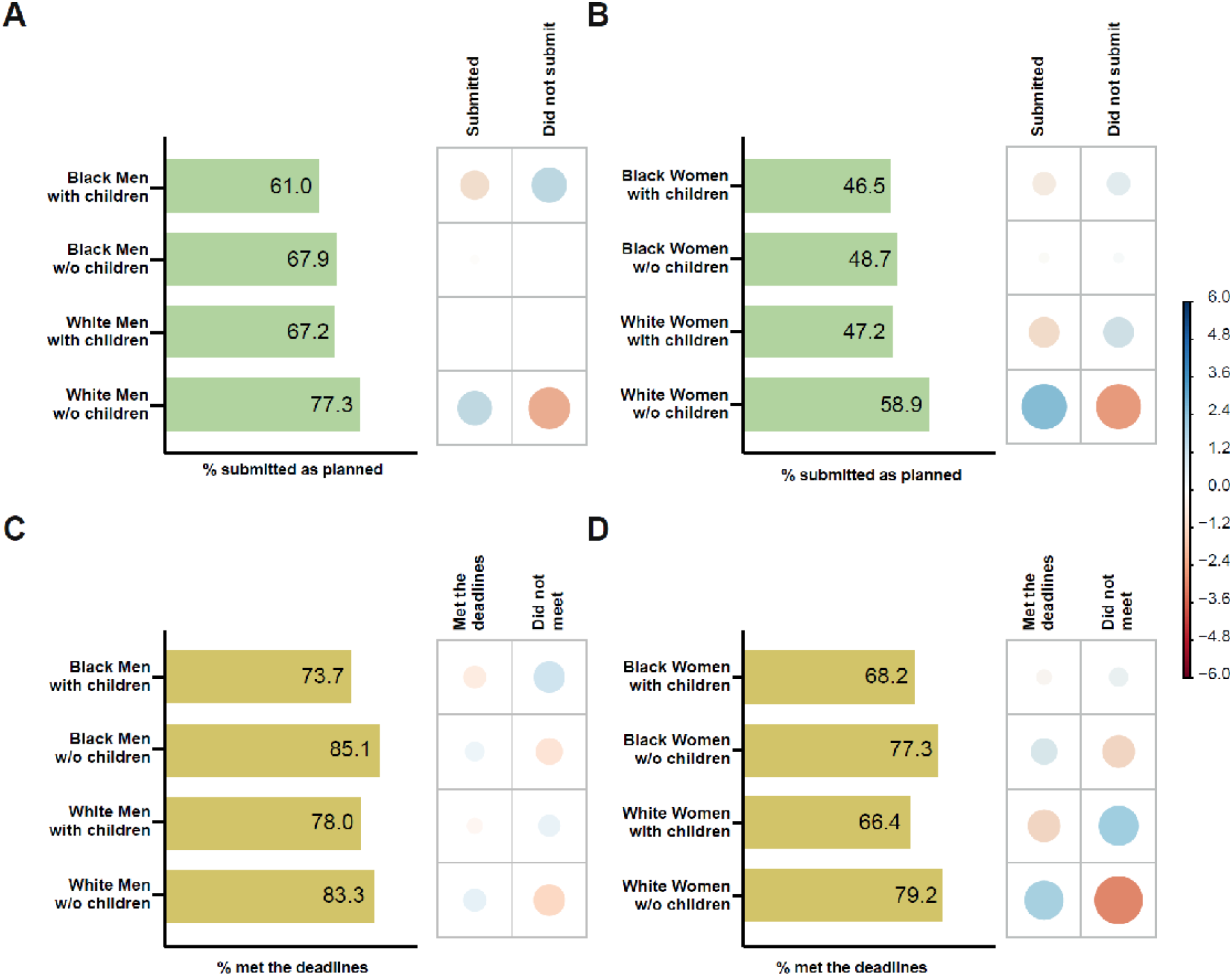
The influence of race and parenting on the manuscript submissions (A, B) and meeting the deadlines (C, D) during COVID-19 pandemic. For each figure, the graph on the left-hand side represents the percentage of respondents that submitted manuscripts as planned (A, B) or met the deadlines (C, D), while on the right-hand side the correlation plots show Pearson’s chi-squared standardized residuals calculated for each group. Positive residuals (blue) indicate a positive correlation, whereas negative residuals (red) indicate a negative correlation. The size of the circle is proportional to the amount of the cell contribution to the χ2 score. A. Effect of race vs parenting for men on submissions. B. Effect of race vs parenting for women on submissions. C. Effect of race vs parenting for men on meeting the deadlines. D. Effect of race vs parenting for women on meeting the deadlines.

Children age also influenced the productivity. There was a significant difference between men and women depending on the age of the youngest child for the submission of manuscripts (χ2 = 147.95, P < 0.0001) (Fig. 4A). There was a negative association between women with the youngest child ranging from 1 to 6 years-old and submission of manuscripts. The proportion of this group’s submissions is lower than that observed for men with children at the same age (P < 0.001, Bonferroni PostHoc test) (Fig. 4A). Also, the proportion of submissions observed for men with the youngest child age ranging from 7 to 12, 13 to 18, and more than 18 years-old are higher than those observed for women with children for the same ages (P < 0.001 for all comparisons, Bonferroni PostHoc test) (Fig. 4A). There was a significant difference between men and women depending on the age of the youngest child for meeting the deadlines (χ2 = 83.37, P < 0.0001) (Fig. 4B). The proportion of women with the youngest child ranging from 1 to 6, 7 to 12, 13 to 18, and more than 18 years-old that met the deadlines are lower than that observed for men with children at the same ages (P < 0.01 for all comparisons, Bonferroni PostHoc test) (Fig. 4B).

**Figure 4.**
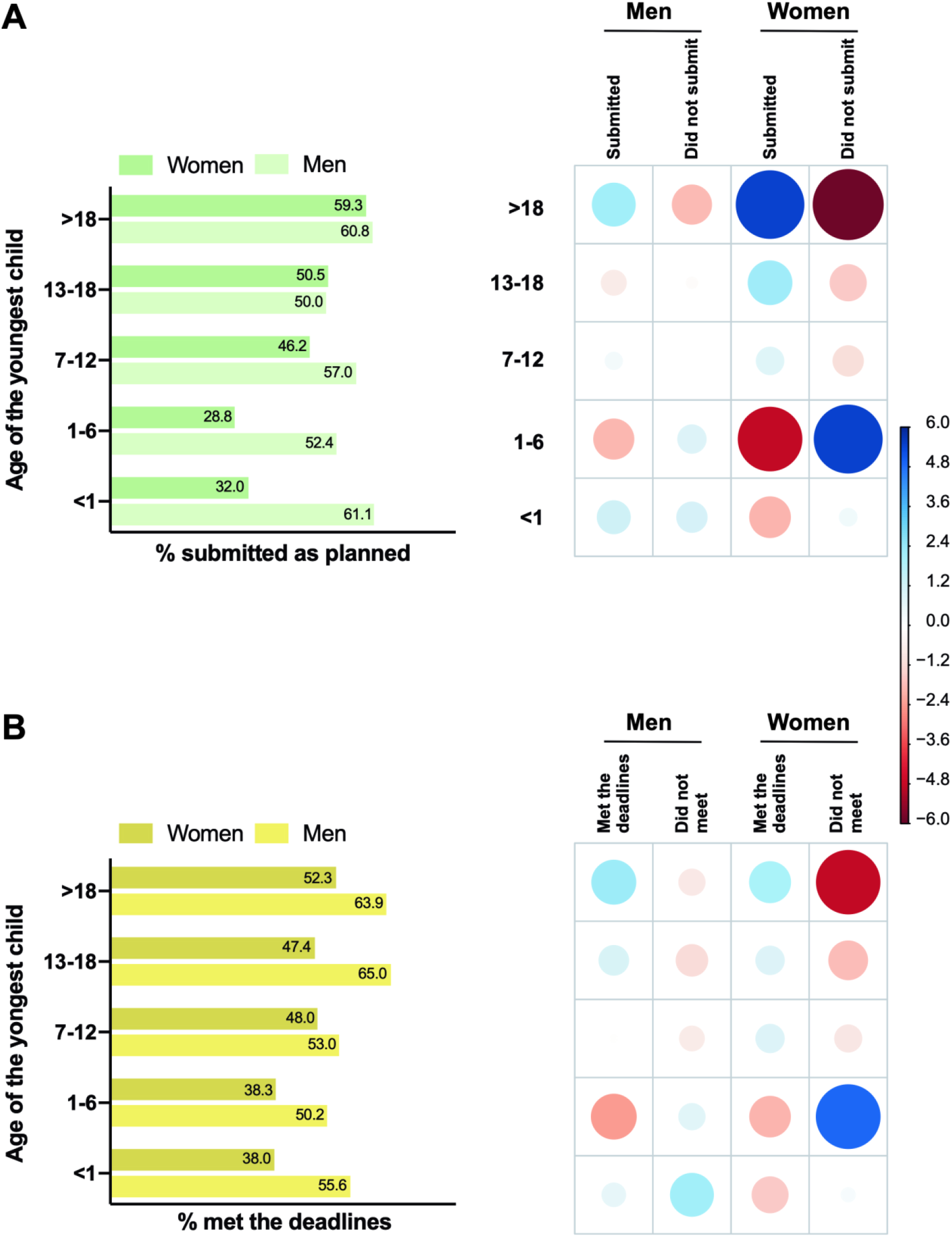
The influence of the youngest child age on the submission of manuscripts as planned (A) and on meeting deadlines (B) during COVID-19 pandemic. The graph on the left-hand side represents the percentage of respondents that submitted manuscripts as planned (A) or met the deadlines (B), while on the right-hand side the correlation plots show Pearson’s chi-squared standardized residuals calculated for each group. Positive residuals (blue) indicate a positive correlation, whereas negative residuals (red) indicate a negative correlation. The size of the circle is proportional to the amount of the cell contribution to the χ2 score. A. Effect of the child age on submission of manuscripts. B. Effect of the child age on meeting the deadlines.

## Discussion

Our findings revealed that male academics productivity has been less affected by the pandemic circumstances than that of women: they were those most able to submit manuscripts as planned during the pandemic period (Fig. 1A). Parenthood had a strong influence on the ability to submit papers as planned and to meet deadlines during the pandemic period, affecting women with children the most, men without children the least (Fig. 1, C and D). While no evidence was found to an overall racial effect in productivity during the pandemic period (Fig S1), an intersectional look, between the factors gender and race, allowed for the identification of a low effect on the productivity of White men (Fig. 2). Moreover, the intersection between parenthood and race presented a very low impact for male academics, regardless of race, presenting a slight negative effect in their ability to submit papers (Fig. 3A), but not to meet deadlines (Fig. 3B). As for female academics, the factor race presented a strong effect. From an intersectional standpoint, parenthood did not affect the productivity of Black women, whereas it strongly affected the productivity of White women (Fig. 3, C and D). Amongst female academics, the productivity of White women without children was the least affected. We also investigate whether children’s age plays a role in impacting productivity. We found that the younger the children, the greater the impact on women’s productivity both in terms of manuscript submission and the ability to meet deadlines, while the effects on men’s productivity are much lower (Fig. 4A and B). Our results suggest that the factors gender, parenthood and race are impacting the ability to submit manuscripts and to meet deadlines during the pandemic period. Nevertheless, not all scientists are being impacted in the same way: White academic mothers and Black female academics, regardless of motherhood, are the groups taking the strongest hit.

Our study is the first one to provide conclusive data on the forces driving productivity imbalance in science during the pandemic: race and motherhood. Our results for the Brazil scenario echo studies based on the US context that showed that working mothers, including academic ones, might be disproportionately affected by the Covid-19 crisis^8,9^. This exacerbated disparity during the pandemic reflects the historical inequality between the careers of men and women. Women can suffer a decrease in working productivity after the birth of their children^10^. As a result, an increase in the gender gap after motherhood thus occurs in many working areas^10–13^, including academia, where mothers spend 8.5 more hours per week on parenting or domestic tasks, and less time on research than fathers^4^. This asymmetrical division of parenting and domestic tasks can be reflected in a decrease in the number of annual scientific publications^14^, thus affecting the career progression of academic mothers. Interestingly, but not surprisingly, children’s age had an impact on mothers’ productivity. Young children require much more attention and care, in addition to demands related to having time to homeschool children during the pandemic. Unfortunately, gender inequality intersects with the racial profile of academics. Indeed, Black women are greatly underrepresented in science, particularly in STEM fields^15^: in the US, for instance, only 2% of practicing scientists and engineers are Black women^16^. They represent only 2% of full-time professors in research institutions^7^. In Brazil, Black women are also underrepresented in science representing only 3% of PhD supervisors^17,18^. Our data confirm this excluding scenario, by showing that Black female academics, regardless of the motherhood factor, are the most affected group by pandemic circumstances. Interestingly, the productivity effect on Black women without children differs from that of White women without children. This evidence reveals an important influence of the race factor in science. The reasons behind it are still debatable, but systemic racism, lack of representation, race-based stereotypes, low socio-economics conditions should be particularly regarded as probable reasons^15^. This is a huge issue because diversity is a keystone for building a better and innovative science^19^.

In summary, our findings revealed that female academics, especially Black female ones, and mothers (regardless of race), are paying most of the pandemic impacts bill. This fact could lead to an unprecedented increase in both gender and race gaps in science. Our study strongly recommends the implementation of policies and actions in order to mitigate this reality. The international academic community needs flexibility in institutional policies from research institutions and funding agencies, such as the postponement of deadlines for grant proposals and reports. Furthermore, funding agencies should consider creating grants designed to benefit Black scientists and academics with families. In times of growing compassion, we invite the entire scientific community to make science more diverse and fairer after the pandemic.

## Methods

This project was approved by the Ethics Committee of the Federal University of Rio Grande do Sul (CAAE 82423618.2.0000.5347). This study was conducted by the Parent in Science Movement, which was founded in 2017 in Brazil to fight against the broadly perceived idea that a successful academic career in science is incompatible with having a family, and to provide support to mothers facing this conflict in their lives. The movement main goal is to promote institutional changes to ensure women are not forced to choose between motherhood and a career in science. The study was performed using an online research survey that was available on the web between April 22^nd^ and May 25^th^, 2020. In this period, Brazilian day cares, schools, and universities were closed due to the COVID-19 pandemic. This survey was posted in social media and was e-mailed to universities and research centers based in Brazil. The snowball sampling technique was also used, where existing study subjects recruit future subjects from among their acquaintances. The survey took approximately 5 minutes to complete. Participants who failed to fully complete the questionnaire were excluded. The final sample was composed of 3,345 individuals, predominantly women (68.4 %). Higher rates of female respondents in studies targeting university faculty members have been previously reported^20^.

The psychometric questionnaire was specially developed to assess the impact of social distancing, due to COVID-19, on the productivity of researchers of both sexes with and without children. It consisted of 25 questions, including information about the researcher’ demographics (country region, gender, and race), work setting, children, and remote work conditions (see a complete version of the questionnaire in the supplementary material). Productivity was assessed by the researchers ability of submitting papers and meeting deadlines during the social distancing period.

Data are presented as the percentage of respondents that were able to submit papers as planned and to meet deadlines related to grants/fellowships proposals and/or project/funding reports within each analyzed group. Statistical analysis to test for differences between groups (men and women, with or without children, also stratified by the age of the youngest child, from different race/ethnicity) was performed using a chi squared test. Chi squared analysis was performed in R version 4.0.0 using the chisq.test function. Pearson residuals plots were generated with the corrplot package (version 0.84). Finally, pair comparison between groups from statistically significant Chi-squared tests were run with the chisq.multcomp function of the RVAideMemoire package (version 0.9 - 77), using Bonferroni correction of p-values. Significance level was set at 0.05.

**Table 1.**
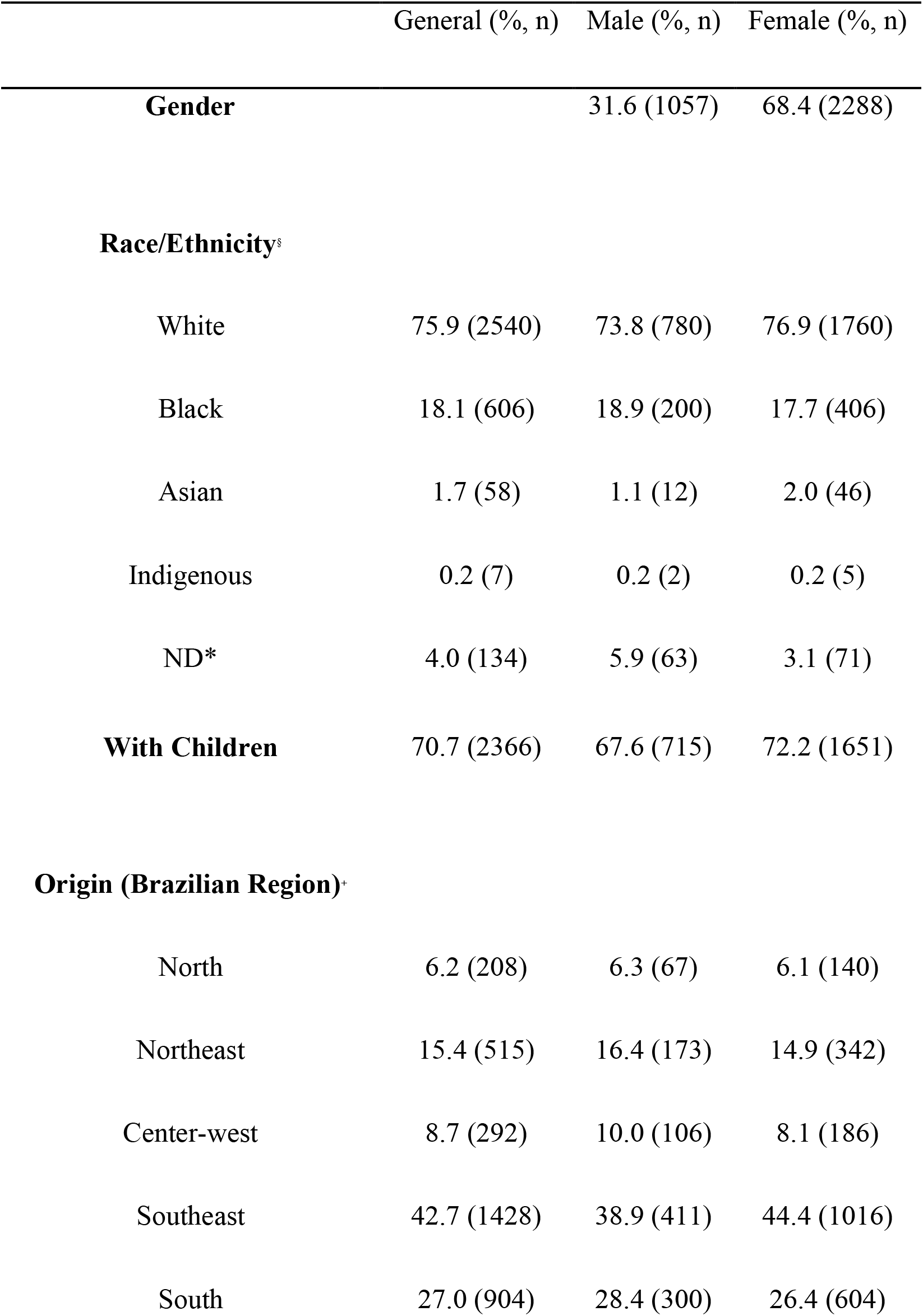
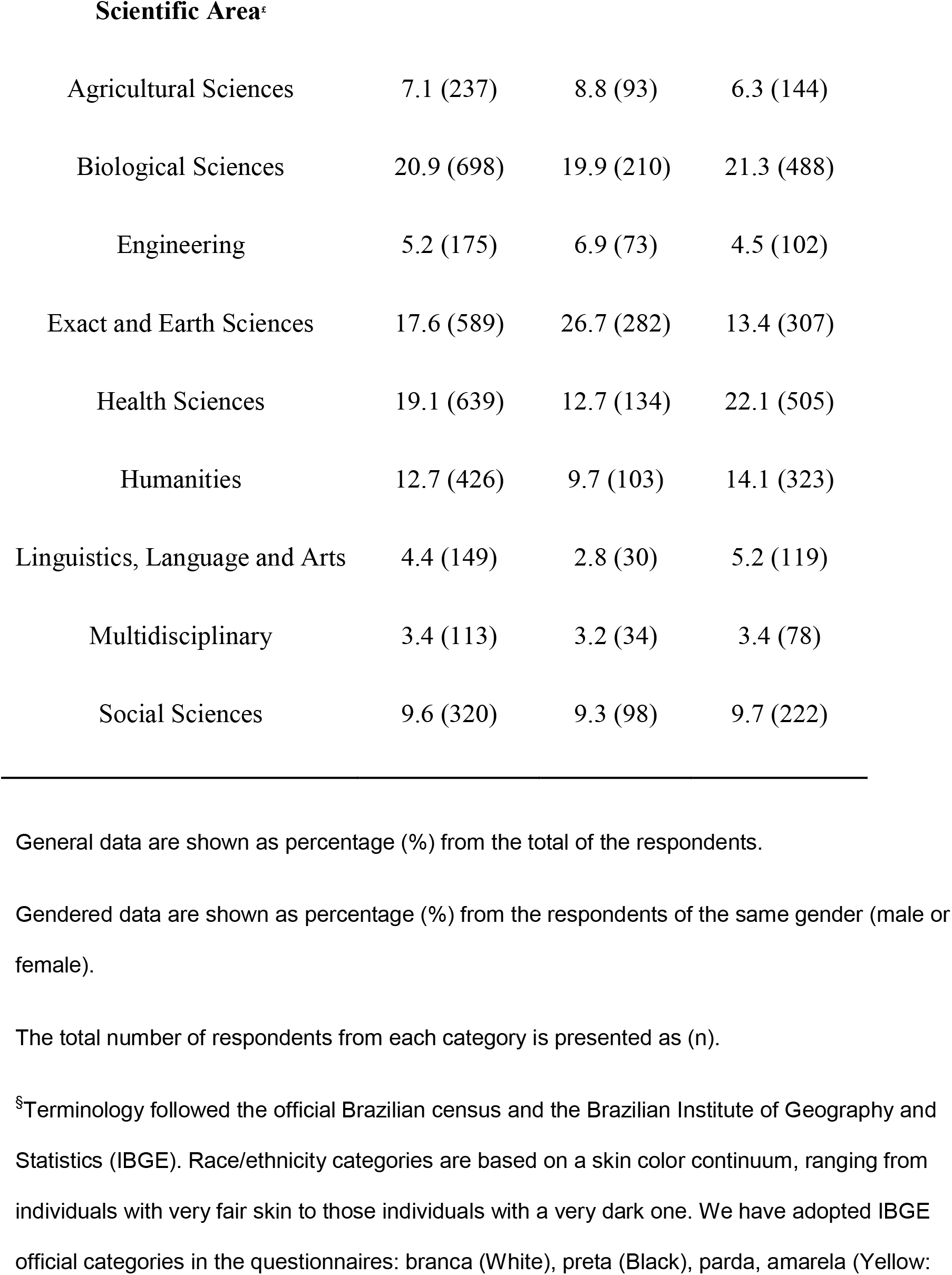
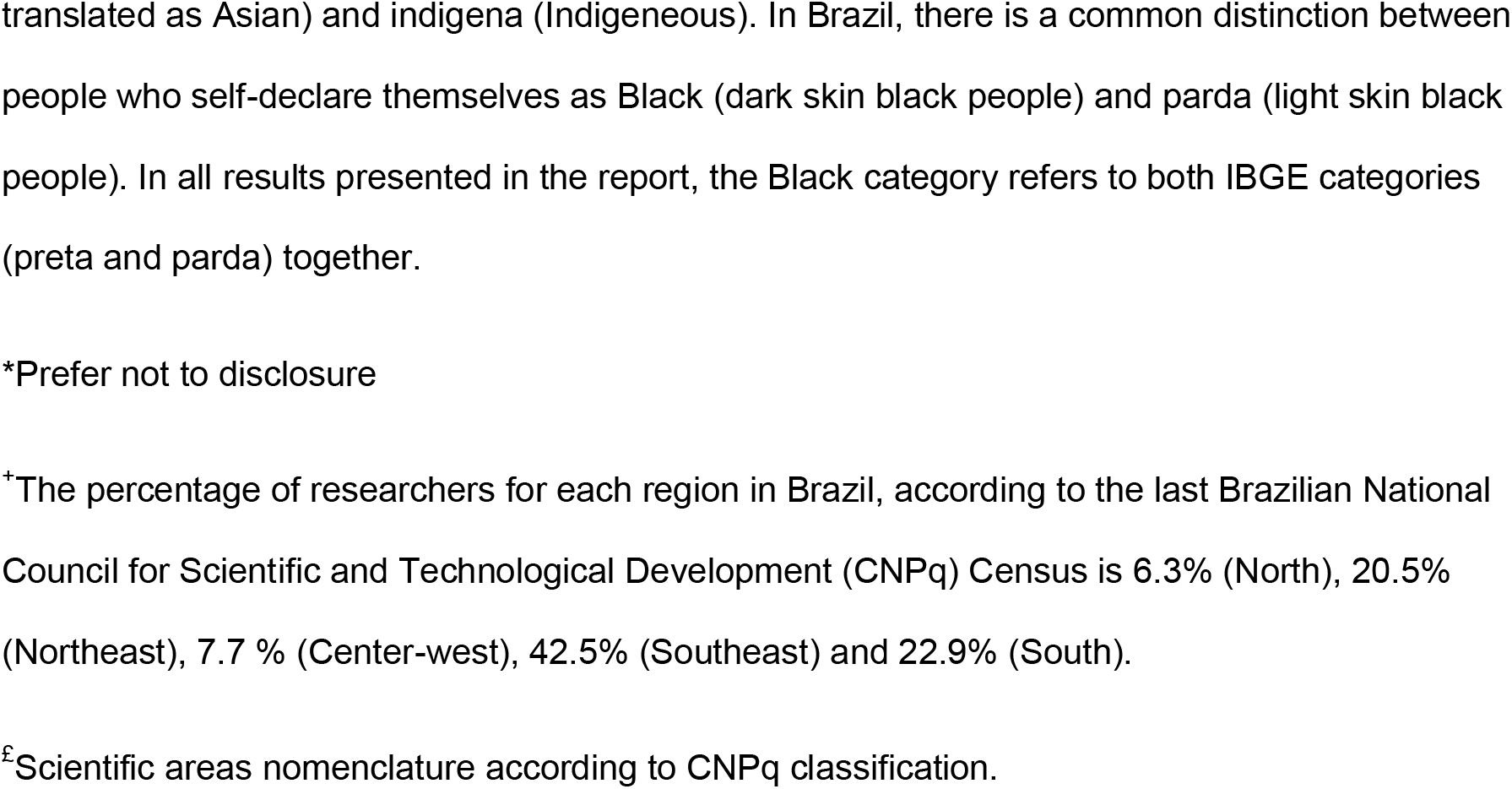
Characterization of the sample included in the study (3,345 respondents).

|  | General (% , n) | Male (% , n) | Female (% , n) |
| --- | --- | --- | --- |
| **Gender** |  | 31.6 (1057) | 68.4 (2288) |
| **Race/Ethnicity<sup>s</sup>** |  |  |  |
| White | 75.9 (2540) | 73.8 (780) | 76.9 (1760) |
| Black | 18.1 (606) | 18.9 (200) | 17.7 (406) |
| Asian | 1.7 (58) | 1.1 (12) | 2.0 (46) |
| Indigenous | 0.2 (7) | 0.2 (2) | 0.2 (5) |
| ND* | 4.0 (134) | 5.9 (63) | 3.1 (71) |
| **With Children** | 70.7 (2366) | 67.6 (715) | 72.2 (1651) |
| **Origin (Brazilian Region)\*** |  |  |  |
| North | 6.2 (208) | 6.3 (67) | 6.1 (140) |
| Northeast | 15.4 (515) | 16.4 (173) | 14.9 (342) |
| Center-west | 8.7 (292) | 10.0 (106) | 8.1 (186) |
| Southeast | 42.7 (1428) | 38.9 (411) | 44.4 (1016) |
| South | 27.0 (904) | 28.4 (300) | 26.4 (604) |

| **Scientific Area<sup>§</sup>** |  |  |  |
| --- | --- | --- | --- |
| Agricultural Sciences | 7.1 (237) | 8.8 (93) | 6.3 (144) |
| Biological Sciences | 20.9 (698) | 19.9 (210) | 21.3 (488) |
| Engineering | 5.2 (175) | 6.9 (73) | 4.5 (102) |
| Exact and Earth Sciences | 17.6 (589) | 26.7 (282) | 13.4 (307) |
| Health Sciences | 19.1 (639) | 12.7 (134) | 22.1 (505) |
| Humanities | 12.7 (426) | 9.7 (103) | 14.1 (323) |
| Linguistics, Language and Arts | 4.4 (149) | 2.8 (30) | 5.2 (119) |
| Multidisciplinary | 3.4 (113) | 3.2 (34) | 3.4 (78) |
| Social Sciences | 9.6 (320) | 9.3 (98) | 9.7 (222) |

General data are shown as percentage (%) from the total of the respondents.
Gendered data are shown as percentage (%) from the respondents of the same gender (male or female).
The total number of respondents from each category is presented as (n).
<sup>§</sup>Terminology followed the official Brazilian census and the Brazilian Institute of Geography and Statistics (IBGE). Race/ethnicity categories are based on a skin color continuum, ranging from individuals with very fair skin to those individuals with a very dark one. We have adopted IBGE official categories in the questionnaires: branca (White), preta (Black), parda, amarela (Yellow:
translated as Asian) and indigena (Indigeneous). In Brazil, there is a common distinction between people who self-declare themselves as Black (dark skin black people) and parda (light skin black people). In all results presented in the report, the Black category refers to both IBGE categories (preta and parda) together.
\*Prefer not to disclosure
<sup>+</sup>The percentage of researchers for each region in Brazil, according to the last Brazilian National Council for Scientific and Technological Development (CNPq) Census is 6.3% (North), 20.5% (Northeast), 7.7 % (Center-west), 42.5% (Southeast) and 22.9% (South).
<sup>£</sup>Scientific areas nomenclature according to CNPq classification.

## Acknowledgments

We would like to acknowledge our children who are the reason we keep on the fight for a fairer world. Also, we are thankful to all scientists that answered our survey. This work was supported by the Conselho Nacional de Desenvolvimento Científico e Tecnológico (CNPQ), and the Fundação de Amparo à Pesquisa do Estado do Rio de Janeiro (FAPERJ).

## Author Contributions

Conceptualization - FS, LK, EZ, FR, RCS, AN, IVDS, PBMC, ASKT, FPW, FKR, CI, AS, LO. Data curation - FS, LK, EZ, FR, RCS, ZMCL, EL, AN, IVDS, PBMC, ASKT, FPW, FKR, CI, AS. Formal Analysis- FS, LK, EZ, FR, RCS, EL, AN, IVDS, PBMC, ASKT, FPW, FKR, CI, AS, CCS, LO. Funding acquisition - EZ, IVDS, PBMC, CCS, LO. Investigation- FS, LK, EZ, FR, RCS, ZMCL, EL, AN, IVDS, PBMC, ASKT, FPW, FKR, CI, AS, CCS, LO. Methodology - FS, LK, EZ, FR, RCS, ZMCL, EL, AN, IVDS, PBMC, ASKT, FPW, FKR, CI, AS, CCS, LO. Project administration - FS, LK, EZ, FR, RCS, ZMCL, EL, AN, IVDS, PBM, ASKT, FPW, FKR, CI, AS, CCS, LO. Resources – LK, CCS. Supervision - FS, LK, EZ, FR, RCS, LO. Visualization - FS, LK, EZ, FR, RCS, EL, AN, PBM, ASKT, FPW, FKR, CCS, LO. Writing – original draft FS, LK, EZ, FR, RCS, ZMCL, EL, AN, IVDS, PBM, ASKT, FPW, FKR, CI, LO. Writing – review & editing - FS, LK, EZ, FR, RCS, ZMCL, EL, AN, IVDS, PBM, ASKT, FPW, FKR, CI, AS, CCS, LO.

