## Supplementary information for "Gender, race and parenthood impact academic productivity during the COVID-19 pandemic: from survey to action"

### Supplementary Results

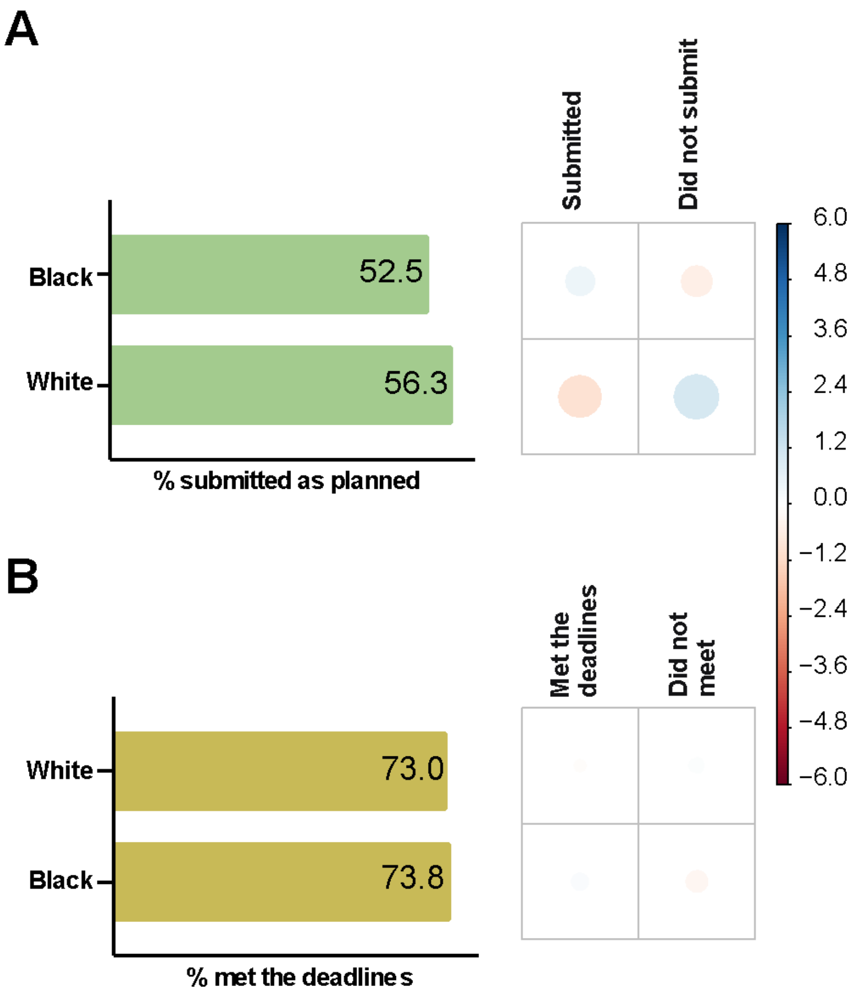


**Supplementary Figure 1.** The influence of race in manuscript submission and meeting deadlines during COVID-19 pandemic. The graph on the left-hand side represents the percentage of respondents that submitted manuscripts as planned (A) or met the deadlines (B), while on the right-hand side the correlation plots show Pearson's chi-squared standardized residuals calculated for each group. Positive residuals (blue) indicate a positive correlation, whereas negative residuals (red) indicate a negative correlation. The size of the circle is proportional to the amount of the cell contribution to the $\chi 2$ score. A. Race effect on submissions. B. Race effect on meeting deadlines.

**Data S1. The complete version of the questionnaire**

Scientists, productivity and the pandemic: which are the impacts?

*Required

What is your current institutional position?*

☐Substitute Professor

☐Adjunct Professor

☐Associated Professor

☐Full Professor

☐Researcher (non-teaching)

☐Other:

How long have you been employed as a Professor/Researcher (do not count time spent as a graduate student/post-doc)*

☐Under a year

☐1 to 5 years

☐5 to 10 years

☐10 to 15 years

☐Over 15 years

You are employed by a _______ institution*

☐Public

☐Private

In which region is your institution located?*

☐North

☐Northeast

☐Centerwest

☐Southeast

☐South

Which institution are you affiliated to? Please write the name of the institution, followed by the department. Ex: Federal University of Rio Grande do Sul – Department of Physiology

Your answer

What is your area of knowledge?*

☐Agricultural Sciences

☐Biological Sciences

☐Health Sciences

☐Humanities

☐Social Sciences

☐Linguistics, Language and Arts

☐Exact and Earth Sciences

☐Engineering

☐Multidisciplinary

Do you have a CNPq productivity fellowshipA?*

☐No

☐Senior

☐1A

☐1B

☐1C

☐1D

☐2

How do you identify yourself?*

☐Male

☐Female

☐Other:

According to the IBGE census race and color categories, you declare to be:*

☐Indigenous

☐Asian

☐Black

☐Parda

☐White

☐I do not wish to answer

When did your institution suspend in-person activities? Please answer using the DD/MM/2020 format (in case they have not been suspended, please answer "not suspended)*

Your answer

Which situations best define how your professional life is being impacted by the COVID-19 pandemic? Please mark all relevant alternatives.*

☐I have not experienced any impact in my professional life until now.

☐I can dedicate more time to research, as I have fewer demands for other activities (for example, in-person classes and non-mandatory meetings).

☐My workplace is closed, but research activities are fully continuing remotely.

☐My workplace is closed, but research activities are continuing remotely, partially.

☐My workplace is closed, and I cannot perform research activities remotely.

☐The laboratory where I work continues to function only for essential activities (management of animal facilities/herbarium/cell culture, etc.)

☐My group has suspended fieldwork for all research projects.

☐My group has substituted fieldwork with activities that can be performed remotely.

Are you developing on-line teaching activities, substituting in-person classes?*

☐I am not teaching at the moment

☐Yes - fully

☐Yes - partially

☐No

Do you coordinate any outreach activities? *

☐Yes

☐No

In case you answered ‘yes’ in the previous question, check the alternative that best defines your activity status:

☐Outreach activities continue normally.

☐Outreach activities have been fully suspended.

☐Outreach activities have been partially suspended.

In your opinion, is this period of closed institutions and imposed adaptation to remote work affecting your productivity?*

☐No

☐Yes – Positively

☐Yes – Negatively

So far, has the pandemic situation impacted how you meet deadlines?*

☐No impact, since I was able to perform necessary tasks

☐No impact, as the funding agency/institution delayed deadlines due to the pandemic

☐I lost deadlines for grant/fellowship applications and/or research/financial report submission

Regarding paper submissions, you:*

☐Have not tried to submit any papers during the social distancing period.

☐Could not finish a paper for submission during the social distancing period.

☐Submitted papers as planned.

How many Master students do you currently advise or co-advise? Answer with numbers only

Your answer

How many PhD students do you currently advise or co-advise? Answer with numbers only

Your answer

Regarding the graduate students you advise, have there been changes in the scheduled defense date for at least one student?

☐ No

☐ Yes, for at least one scheduled defense in the period from March 19 to May 17, in accordance with recommendations in CAPES ordinance nº 36 of 2020.

☐ Yes, for at least one scheduled defense in the first semester of 2020, but outside the expected period in accordance with CAPES ordinance nº 36 of 2020.

☐ Yes, for at least one scheduled defense in the second semester of 2020.

Do you have children?*

☐ No

☐ Yes - one

☐ Yes - two

☐ Yes - three

☐Yes - four or more

What is the age of your youngest child? Answer with numbers only. If younger than one, answer "0".

Your answer

In case you have more than one child, inform their age (numbers only) leaving a space between each number. If younger than one, answer "0".

Your answer

How would you describe your support network DURING THE PANDEMIC PERIOD? Choose the option that best describes your situation.

☐I am sharing childcare with my partner

☐I have a support network – person/people other than my partner

☐I do not have a support network – my partner maintained work activities at his workplace

☐I do not have a support network

☐Other

Are there any factors in your current situation that impact your remote work? Check all correct alternatives*

☐Yes – caring for child(ren) routine

☐Yes – caring for child(ren) homework

☐Yes - caring for child(ren) with disabilities

☐Yes - caring for elderly relatives or other family members (not children)

☐Yes - caring for home/house duties

☐No - I am able to work remotely in a comparable way to the period before the pandemic.

☐Other:

Please, leave your comments here.

Your answer

Notes:

A CNPQ (Conselho Nacional de Desenvolvimento Científico e Tecnológico) is one of the major science funding agencies in Brazil. The Productivity Fellowship is granted to researchers that meet specific criteria and are highly productive in their areas. Evaluation committees receive individual applications and rank scientists depending on these criteria. The ranking in increasing order starts in 2 up to 1A.
